# 3D Ultrastructure of the Cochlear Outer Hair Cell Lateral Wall Revealed By Electron Tomography

**DOI:** 10.1101/534222

**Authors:** William Jeffrey Triffo, Hildur Palsdottir, David Gene Morgan, Kent L. McDonald, Robert M. Raphael, Manfred Auer

## Abstract

Outer hair cells in the mammalian cochlea display a unique type of voltage-induced mechanical movement, termed electromotility, which amplifies auditory signals and contributes to the sensitivity and frequency selectivity of mammalian hearing. Electromotility occurs in the outer hair cell (OHC) lateral wall, and it is not fully understood how the supramolecular architecture of the lateral wall enables this unique form of cellular motility. Employing electron tomography of high-pressure frozen and freeze-substituted OHCs, we visualized the 3D structure and organization of the membrane and cytoskeletal components of the OHC lateral wall. The subsurface cisterna (SSC) is a highly prominent feature, and we report that the SSC membranes and lumen possess hexagonally ordered arrays of particles that endow the SSC with a previously unrealized anisotropic structural rigidity. We also find the SSC is tightly connected to adjacent actin filaments by short filamentous protein connections spaced at regular intervals. Pillar proteins that join the plasma membrane to the cytoskeleton appear as variable structures considerably thinner than actin filaments and significantly more flexible than actin-SSC links. The structurally rich organization and rigidity of the SSC coupled with apparently weaker mechanical connections between the plasma membrane and cytoskeleton reveal that the membrane-cytoskeletal architecture of the OHC lateral wall is more complex than previously appreciated. These observations are important for our understanding of OHC mechanics and need to be considered in computational models of OHC electromotility that incorporate subcellular features.

## Introduction

Sound is processed in the mammalian cochlea by two distinct types of auditory hair cells located in the organ of Corti: inner hair cells and outer hair cells. Inner hair cells transduce sound-induced vibrations into neural signals, while outer hair cells (OHCs) serve to amplify these vibrations and enhance auditory sensitivity and frequency selectivity [1–4]. The three rows of cylindrically-shaped OHCs make a major contribution to the passive mechanics of the organ of Corti and undergo active length changes in response to electrical stimulation [5, 6]. OHC electromotility is elicited by changes in the transmembrane potential that result from hair bundle displacement, with hyperpolarization causing cell elongation, and depolarization causing cell shortening [7, 8].

The process of electromechanical transduction occurs in the lateral wall of the OHC [9, 10], which extends from just below the cuticular plate at the apex of the cell down to the basally located nucleus. Several early studies provided evidence that electromotility is based in the basolateral plasma membrane [11–13], and phenomenological models have been developed to account for experimental observations [1, 14, 15]. Although the discovery of the transmembrane protein prestin [16] provided a molecular basis for a membrane-based electromechanical transduction process [4, 17–19], there are still many unanswered mechanistic questions regarding the relation between prestin activity and cellular length changes. Current models of OHC electromotility posit that a plasma membrane-resident motor generates the force required for cell length changes, either through area-change [20, 21] or nanoscale bending [22]. However, a full biophysical understanding of OHC mechanics and electromotility requires more detailed knowledge of the 3D sub-cellular and molecular architecture of the OHC lateral wall.

The current structural understanding of the outer hair cell lateral wall represents an amalgamation of results from studies spanning multiple decades, utilizing transmission electron microscopy (TEM), scanning electron microscopy (SEM) and more recently atomic force microscopy. The lateral wall can be viewed as a trilaminate composite [23], consisting of the plasma membrane (PM), an underlying cytoskeletal network, and an adjacent system of circumferential lamellar organelles known as the subsurface cisternae (SSC). In freeze-fracture, ~10 nm intramembranous particles were reported in the PM that have been ascribed to the membrane motor protein prestin [13, 24, 25]. The cytoskeletal network lying in the ~30 nm wide extracisternal space (ECS) between the PM and SSC is often referred to as the cortical lattice (CL). It consists of patches of locally parallel, circumferentially oriented filaments of actin connected by axially oriented cross-links, which were assumed to be spectrin polymers [24, 26–29]. This architecture was used to explain observations that the circumferential stiffness of the OHC was greater than its axial stiffness [30]; the resulting anisotropy in lattice stiffness was purported to direct the energy from an isotropic plasma membrane motor down the axis of the cell. A density spanning the ECS between the PM and actin filaments has been observed and called the pillar due to its morphology; estimates on pillar width range from 6-10 nm [24, 29, 31]. In mechanical models of electromotility, the pillar is presumed to provide the necessary coupling of forces generated in the membrane to the underlying cytoskeleton to generate whole-cell shape changes [15, 23].

While the SSC is a prominent sub-structure in the lateral wall, its role in OHC function remains obscure. The SSC closely abuts the cortical lattice (CL), and the number of concentric SSC layers varies according to species, position of the OHC along the cochlea, and longitude along the lateral wall of an individual OHC [24, 32, 33]. One SSC layer occupies similar volume when compared to the PM/CL complex; a previous study reports an average cisternal width of 27 nm [34], comparable to the span of the extracisternal space (ECS). No study to date has shown direct interaction of the SSC with the PM/CL components underlying the motile mechanism, and thus it has been unclear whether the SSC played a role in electromotility.

Prior to this study, the characterization of the OHC lateral wall by electron microscopy techniques has been limited by sample preservation and by the conventional 2D projection imaging of a 3D structure, resulting in superposition of molecules along the electron path. However, a full understanding of electromotility requires an accurate 3D depiction of the dimensions and arrangement of the molecular constituents present within the OHC lateral wall. This is essential for understanding how the micromechanical 3D architecture gives rise to the passive mechanical properties of the OHC, and to assess whether electromechanical force transmission by prestin in the PM alone is likely to be sufficient to deform the entire OHC lateral wall. To address this problem, we employed high-pressure freezing / freeze-substitution [35] to guarantee high fidelity sample preservation, along with electron tomography for 3D imaging of the OHC lateral wall.

Our research has revealed several major new findings. First, both the SSC membrane and its lumenal material are composed of an intricate and highly organized structure not previously reported in any intracellular organelle. Second, the actin filaments of the cortical lattice appear tightly connected to the adjacent SSC membranes by previously unresolved short linkages. In contrast, the pillar proteins that connect the plasma membrane to the cytoskeleton are thinner and significantly more variable than previously observed. Thus, these observations reveal the SSC to be a structurally rich organelle tightly associated with the actin cytoskeleton. Overall, these ultra-structural findings significantly advance our understanding of the subcellular architecture of the OHC lateral wall and provide insight into how this architecture contributes to the overall mechanical properties of the cell.

## Results

### Advantages of High Pressure Freezing / Freeze Substitution and Electron Tomography

Multiple preparation protocols were evaluated using both dissected organ of Corti along with decalcified intact cochleae. These included conventional preparation protocols using paraformaldehyde / glutaraldehyde followed by osmium tetroxide containing potassium ferricyanide, progressive lowering of temperature (PLT) dehydration prior to resin embedding, and high-pressure freezing (HPF) followed by freeze-substitution (FS) prior to resin embedding. All data presented here are derived from HPF/FS applied to rapidly dissected strips of guinea pig OHCs as described previously[36], as such samples displayed the least extraction and best preservation of cytosolic features as judged by cytoskeletal and SSC details. In total, 23 dual-axis tomograms of the OHC lateral wall were recorded and analyzed for this study.

Figure 1 illustrates two representative regions in different orientations that were selected for tilt-series acquisition; the top row is from an axial cut through an OHC, while the bottom row is from an oblique angle closer to a longitudinal section. The top sample was prepared with 20% BSA as a filler during HPF (appearing as dark contrast outside the cell); the bottom sample was prepared using 20% dextran as filler. Dextran was optically transparent after embedding, allowing for easier sample preparation and orientation; however, these samples are more difficult to section. Samples using 10% glycerol as filler yielded optical transparency along with good sectioning properties, but in the case of guinea pig OHCs, the osmotic load resulted in the collapse of the cells. During isolation of OHC strips for HPF, OHCs often come into close proximity with each other; this often allowed data acquisition from regions where two cells’ worth of lateral wall could be imaged in the same tomogram to maximize the area surveyed.

**Figure 1.**
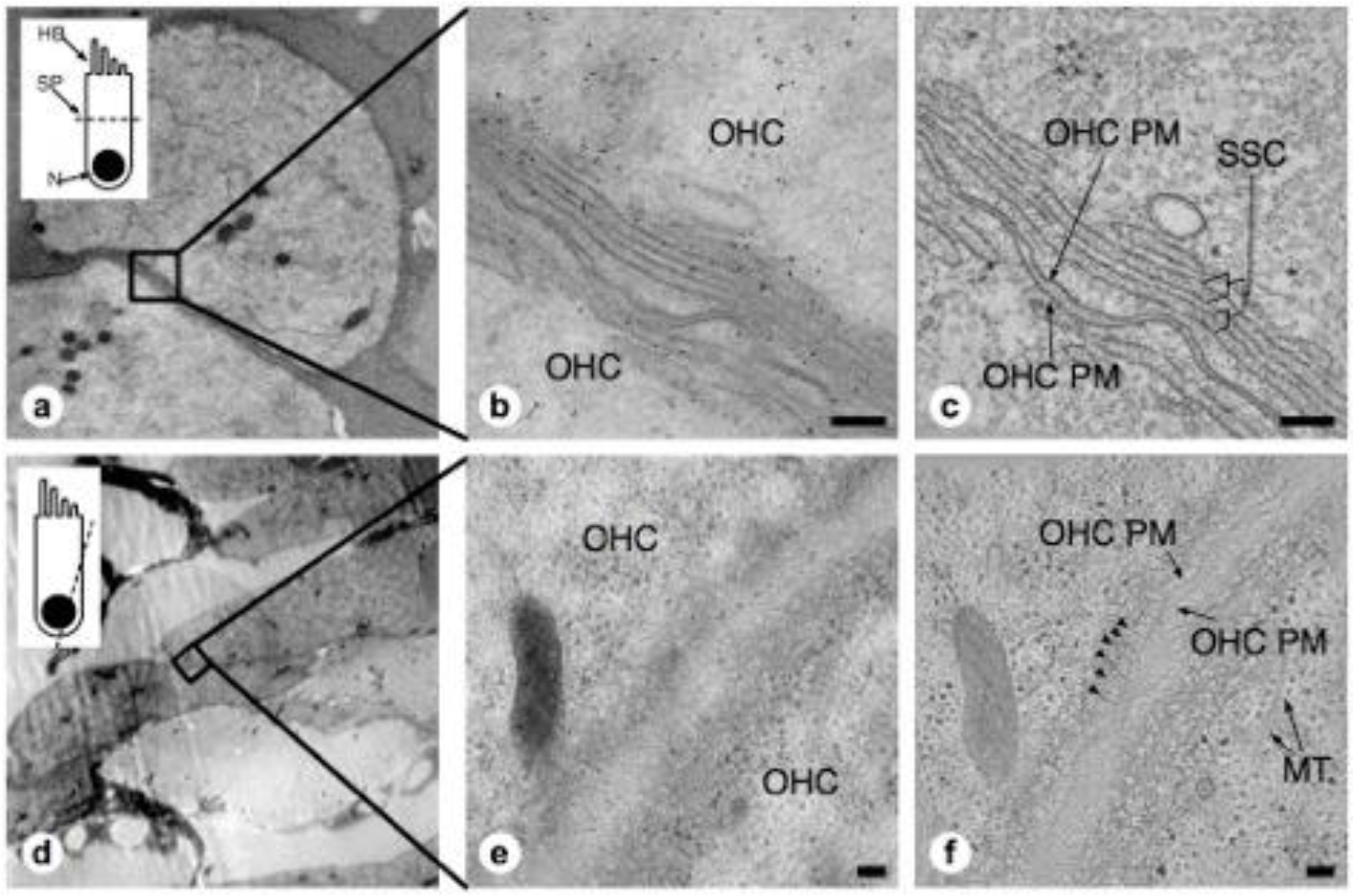
Comparison of projection images and z-planes from OHC lateral wall tomograms. (a,d) low magnification overviews. Inset depicts orientation of plane of sectioning – (a), axial; (d), closer to longitudinal. HB, hair bundle; SP, section plane; N, nucleus. (b,e) projection images from region indicated in black boxes of (a,d). (c,f) mid-volume z-planes from the resulting tomograms. Arrowheads in (f) indicate individual actin filaments in the cortical lattice. MT, microtubules. Scale bars = 100 nm.

The unique advantage of applying electron tomography over conventional 2D projection TEM imaging to the study of OHC lateral wall components is demonstrated by comparison of projection (1b, e) and extracted z-planes from the resulting tomograms (1c, f), revealing the improved resolution in the z-direction of the reconstructions. In panel 1c, internal membranes that delineate the boundaries of each SSC cistern are sharply defined, allowing precise mapping of changes in curvature and continuity in the outermost cistern. In panel 1f, circumferential actin filaments that are nearly parallel to the z-plane are clearly visible just beneath the plasma membrane (marked by arrowheads), whereas in projection views alone these filaments are obscured (1e). This illustrates the advantage of electron tomography compared to projection TEM, along with the challenges of image analysis in a crowded, non-extracted cellular environment.

### Structure of the SSC

In projection, each individual subsurface cisterns appears as a 3-layer “sandwich” consisting of two boundary membranes plus a band of lumenal material (LM), as shown in Figure 2. The LM occupies a third of the lumen volume and lies in between the outer and inner cisternal membranes. This observation was consistent across multiple samples from several freezing sessions with varying HPF filler material concentrations (20% BSA, 20% Dextran, w/v) and freeze-substitution protocols. When arranged in stacks, a sequence of thick and thin bands is routinely observed, corresponding to SSC membrane and LM, respectively (Figure 2a).

**Figure 2.**
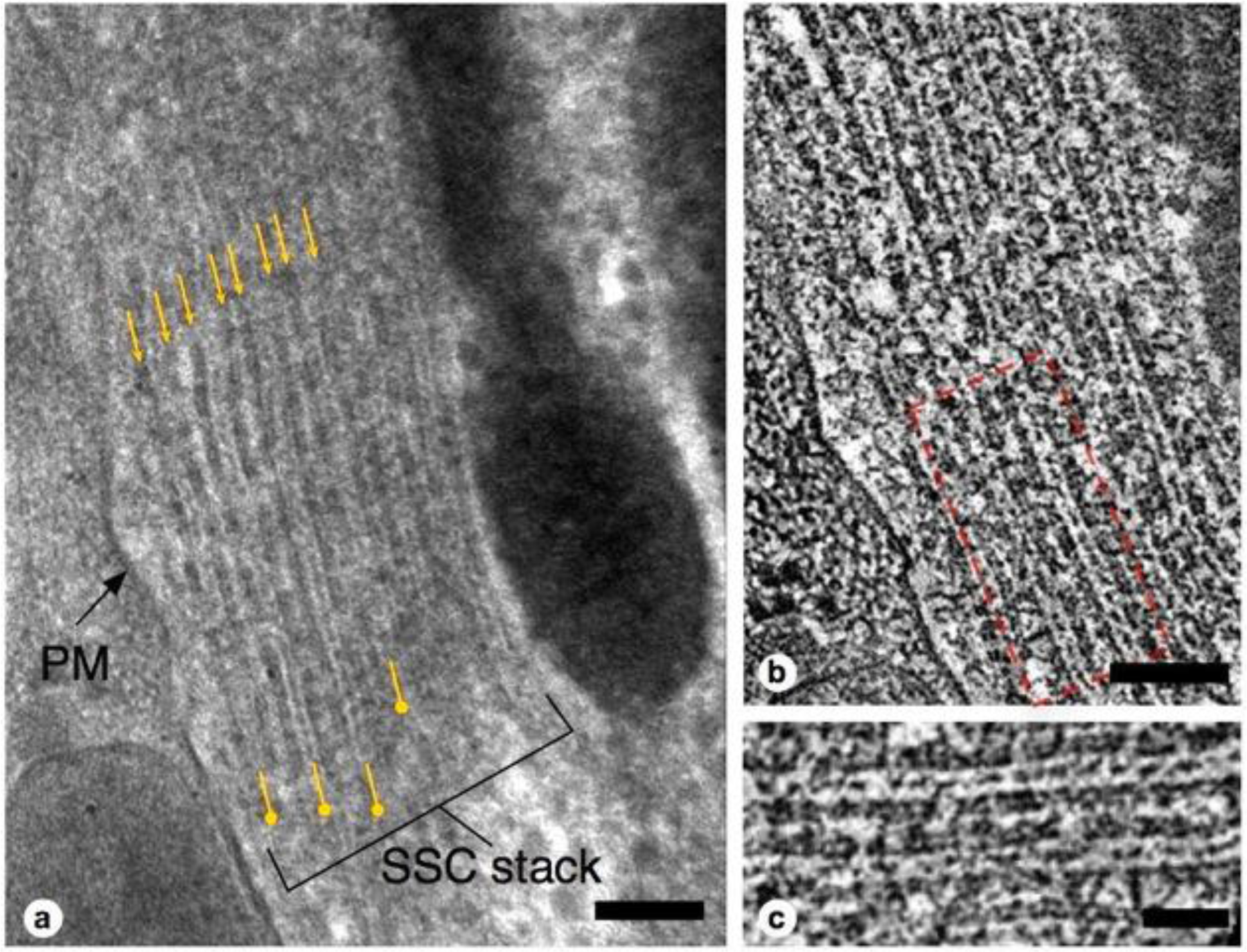
Lateral view of SSC structure. (a) Close-up projection view of lateral wall in longitudinal section. Small arrows indicate membranes of SSC cistern; pushpin annotation denotes lumenal material (LM). (b), 4nm averaged z-plane from tomogram of (a); (c), close-up of two cisterns from red dotted-line box in (b), showing punctate cross-section of LM, occasional contacts between LM and adjacent SSC membrane, and cistern-cistern connections. Scale bar in (a,b) = 100 nm, (c) = 50 nm.

Z-sections from a tomogram of this stack (Figure 2b,c) reveal a quasi-periodic density pattern between adjacent cisternae. Within this region, discrete connections are seen connecting the membranes of neighboring cisternae. In each lumen, the LM appears as a combination of both continuous density and discrete punctae. The individual SSC membranes are of uniform density and appear at least as thick as the plasma membrane (Figure 1c).

A quasi-periodic variation in density is seen within the surface of each SSC membrane, interrupted by sporadic defects in the surface (Figures 3, 4a). No similar pattern is observed in the plasma membrane. The SSC membrane density exhibits a “honeycomb” pattern, with the density arranged along the edges of a hexagonal grid. Considering the local organization to be a hexagonal lattice and measuring along the edges of the resulting equilateral triangles, the average spacing between unit cells is 19.2 +/− 1.9 nm (n = 137). Due to feature size, even gentle curvature of the SSC membrane surface prevents capturing large stretches (more than ~ 100 nm) of the pattern within a single interpolated plane. Thus, surfaces that were modeled using plane-by-plane contours were used to extract volume regions enclosing each SSC membrane for volume rendering. This sequence, along with an en-face rendering of the SSC membrane pattern, is illustrated in Figure 3.

**Figure 3.**
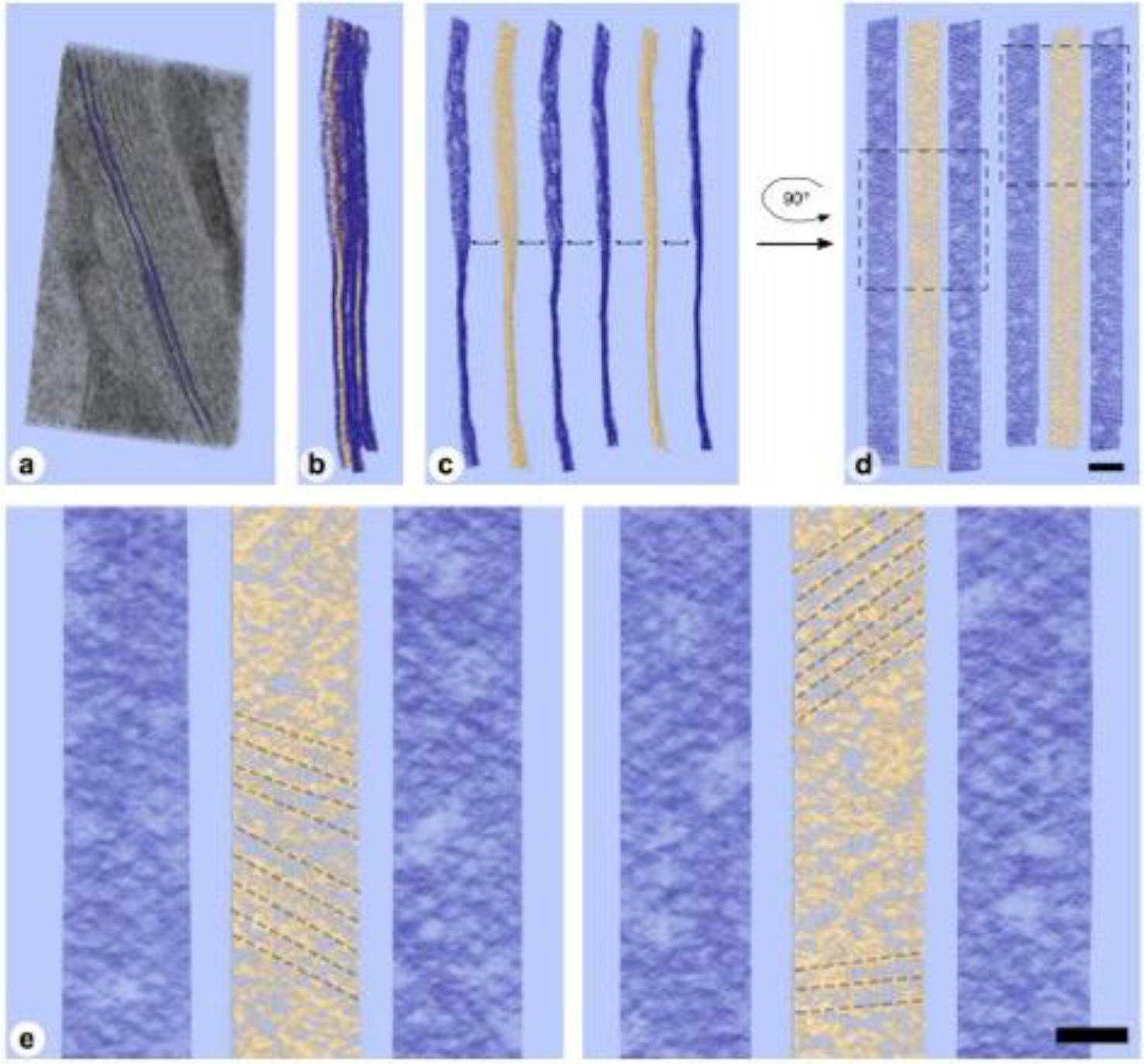
Volume rendering and en-face view of SSC membrane pattern and lumenal material (LM) (a) Volume rendering of tomogram slab with two membranes delineating an SSC cistern rendered as purple surfaces. (b) two cistern density maps extracted from (a) using such surfaces; membrane density is rendered in purple, with LM rendered in gold. (c,d) Separation of membrane and LM along with 90 degree rotation along the y-axis to yield en-face views of the membrane and LM, revealing local hexagonal membrane pattern and patch-like linear organization of LM material. (e) close-up of two regions from (d) indicated by dotted-line boxes. Linear organization of LM, indicated by dotted-line annotation, can also be seen in regions of the volume rendering. The directionality of the LM organization appears to coincide with the lattice lines of the honeycomb pattern in the adjacent SSC membranes. Scale bar in (d) = 100 nm, (e) = 50 nm.

**Figure 4.**
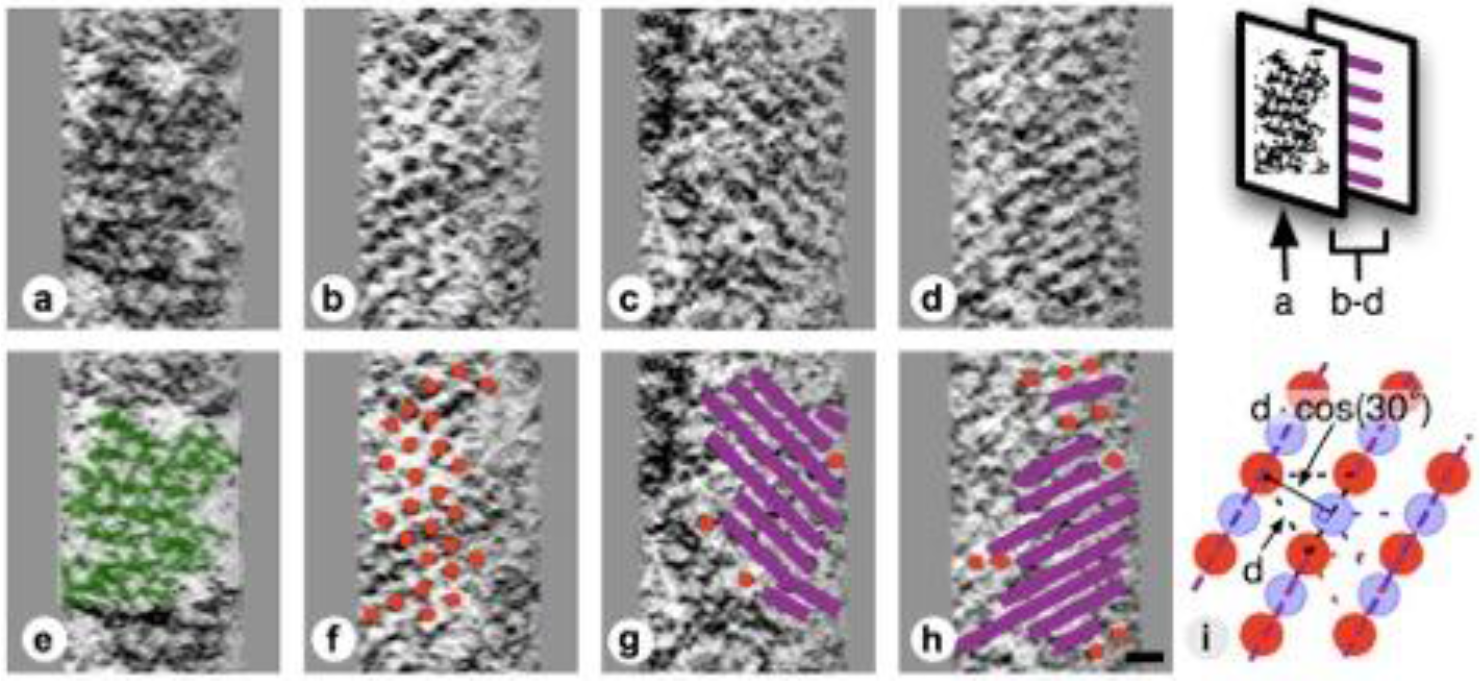
Slice planes through the SSC lumenal material and comparison to local SSC membrane pattern. Images are organized in vertical pairs, with annotation of density superimposed on top of identical bottom image. Diagram at far right indicates location of each pair with respect to cistern. (a,e) SSC honeycomb pattern for reference. (b/f, c/g, d/h) LM is shown to be discrete disks (b/f) that are most commonly seen arranged as continuous bands or filaments (c/g, d/h). (i) Cartoon illustrating how interdigitation of 2 groups of hexagonally arranged disks (red and blue) associated with opposing SSC membranes would give rise to continuous bands of density along lattice lines. Because of the underlying hexagonal organization, the distance between these bands would be related to the hexagonal lattice spacing by cos(30°). Scale bar (h) = 25 nm.

When viewed en-face, individual disk-like punctae, approximately 10 nm in diameter, are seen within the LM. The disks are arranged in a hexagonal pattern with a spacing of 19.1 +/− 2.1 nm (n = 64). The LM is most commonly organized in filament-like parallel rows but discrete quasi-hexagonal patterns are also observed (Figure 4). The row pattern can be interpreted as a quasi-hexagonal appearance, with interdigitating rows of density aligning with the lattice direction of the adjacent SSC pattern (Figure 3d,e). Interdigitating groups of the hexagonally patterned LM disks associated with opposing membranes in an SSC cistern appear to give rise to these continuous bands of density (Figure 4i). In a hexagonal lattice, the orthogonal distance *d* between lattice lines is related to the lattice spacing *a* by the relationship *d* = *a* * cos(30°); for *a* = 19.1 +/− 2.1 nm, *d* = 16.5 +/− 1.8 nm. When organized as rows, the orthogonal distance between adjacent rows is measured to be 16.8 +/− 1.3 nm (n = 61), in excellent agreement with the expected value calculated from the lattice spacing of the individual LM disks. The central LM makes contact with its adjacent SSC membranes (Figure 2b,c), linking together the two membranes of an individual cistern.

### Morphology of the SSC and plasma membrane

In our isolated guinea pig OHCs the outermost layer of the SSC is largely continuous rather than fenestrated, which is often observed by others under conventional preparation methods [32, 34, 37]. The width of each cistern is measured consistently at 28-30 nm. When in layers, the individual cisternae are close together but do not touch, leaving a gap typically less than 20 nm that is bridged by cisternae-cisternae proteinaceous connections. The dextran and BSA fillers used to prevent ice formation during the freezing process contributed ~50 mOsm to the dissecting media, which may explain the undulating appearance of the plasma membrane. Interestingly, the SSC consistently maintains a smooth appearance despite these variations in plasma membrane curvature. This is illustrated in Figure 5a, where the extracted surfaces representing the outermost SSC layer and PM are rendered with PM in light blue, and the SSC membranes in magenta. Occasional discontinuities, or fenestrae, are seen in the outermost SSC layer. These discontinuities are not contained within the thickness of one section, but their boundary indicates a circular, local gap as opposed to a breach around the entire circumference of the cistern.’

**Figure 5.**
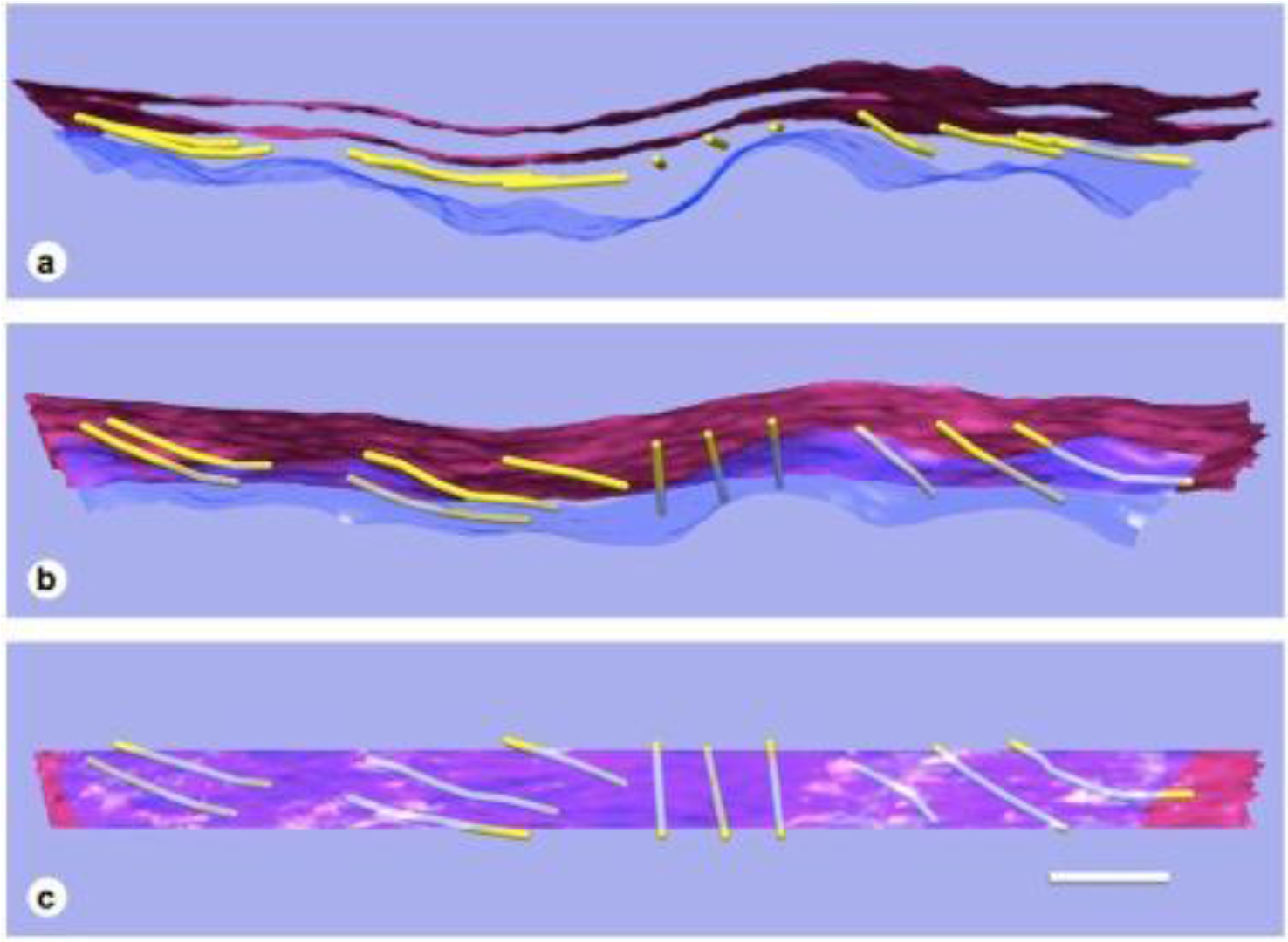
3D relationships of PM, actin, and SSC. Model of single SSC cistern, actin, and PM (a) rotated 45° (b) and 90° (c) about the y-axis. PM in blue / translucent blue; SSC membranes in red, and actin rendered as 8 nm diameter gold filaments. Scalebar = 100 nm.

### SSC is physically connected to Actin

Individual paths of circumferential actin filaments were segmented out and rendered, revealing 8 nm wide cylinders consistent with anticipated actin filament dimensions (Figure 5). The filaments appear to be organized in domains, consistent with the “domain” pattern observed previously [26, 29], with short patches of several filaments running parallel to one another. Furthermore, thin proteinaceous connections between the actin filaments and the PM are also evident, and likely correspond to the “pillar” protein described in previous studies [24, 29, 31] (Figure 6). However, unlike the images depicted in projection studies, pillars were found to be commonly less than half as wide as adjacent actin, with regions of individual pillars frequently as thin as ~3 nm. Of all the components of the lateral wall – PM, CL, SSC, and their associated connections - these pillar connections between PM and actin exhibited the highest variability in morphology. When compared to the adjacent SSC and plasma membranes, the actin filaments maintain a consistent spacing from the SSC membrane even when an undulating plasma membrane is observed (Figure 5). Slice planes through multiple tomographic volumes reveal discrete connections between the actin and SSC membrane, which we refer to as actin-SSC links (Figure 6). There is no previous report of such an actin-SSC linkage, and their existence establishes mechanical coupling between the actin and outermost SSC. Combined with the much thinner than expected pillar connections between the PM and actin, the actin-SSC links complete the physical connection between PM and SSC. It is interesting that in cases of undulating plasma membranes, the pillar structures are found to vary extensively in length whereas the actin-SSC links remain constant.

**Figure 6.**
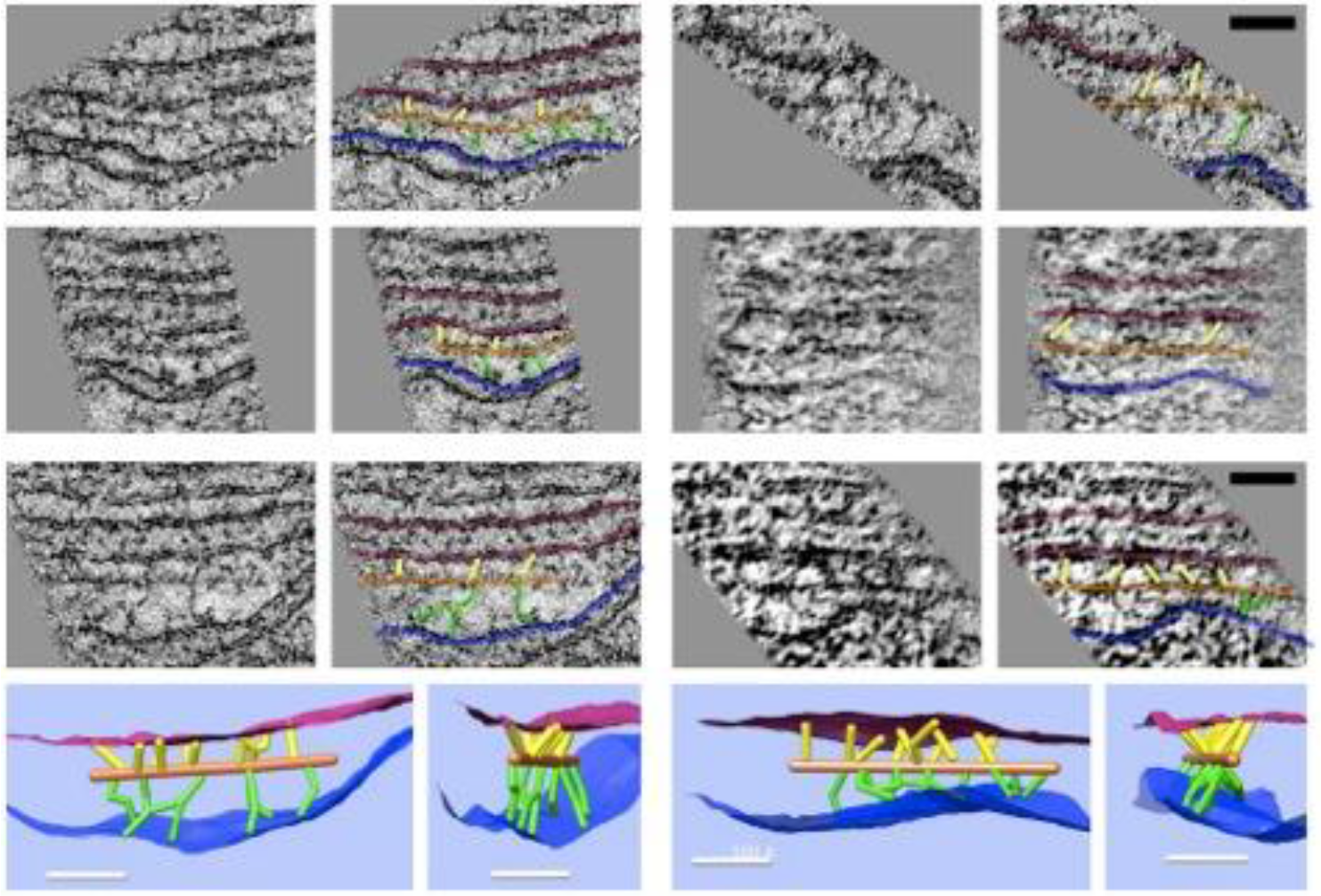
Slice planes and model renderings from tomograms at orientations capturing actin-membrane links. In the top three rows, six separate actin filaments are depicted; each image has an identical copy to its right with overlay in the following colors: red = SSC membrane; blue = PM; orange = actin; green = pillar; yellow = actin-SSC link. In the bottom row, renderings of 3D models corresponding to the filament complexes in the third row are oriented to illustrate the distribution of actin-membrane links not apparent in single 2D slice views. Scalebar = 50 nm.

## Discussion

### SSC membrane and lumen substructure

The hexagonal structure within the membranes of the SSC displays a level of organization not typically associated with the membranes of similar organelles (ER, SER, SR, Golgi apparatus). To our knowledge, this pattern has not been previously described in the SSC membrane, although there is a report that rosette-shaped particles are associated with the lumenal surface of the SSC membrane[38]. However, the proteinaceous pattern we observe is a connected, honeycomb-like structure within the SSC membrane, rather than a field of discrete rosette particles. Therefore, given the repeat length of 19.2 nm and the fact that there is space between the protein densities in the honeycomb-like pattern suggests that such a pattern cannot be explained by dense packing of 10-11 nm intramembranous particles that were described for the PM [13, 24, 25]. In our samples, we applied the same en-face visualization to the PM density, but could not detect any regular protein pattern within the PM.

While generally proteins that do not extend beyond the thickness of the membrane cannot be differentiated from the lipid components of the membrane, we found the SSC membrane pattern to be most prominent under freeze-substitution conditions containing 100% acetone, which are known to deemphasize internal membrane contrast, presumably by destabilization of phospholipid head groups [39]. Therefore, the honeycomb density we have observed in the SSC most likely correlates to an underlying protein scaffold supporting the structure of the membrane.

Knowledge of molecular constituents of the SSC is sparse. In ascribing function to the SSC, correlates have been drawn to similar structures in other motile cells, such as the sarcoplasmic cisternae of striated muscle fibers, with the postulate that the SSC may serve as a calcium store [40, 41]. In this vein, immunogold labeling has provided evidence for the localization of ryanodine receptor (ryR) to the SSC membrane [42]. However, the known dimensions of ryR, with a cytosolic domain 24-25 nm wide [43], are not in agreement with the 19.2 nm repeat we show to be the dominant component of the SSC membrane. Further, ryR has four-fold symmetry [43], which is not compatible with a hexagonal lattice. Thus, the SSC membrane pattern we observe is unlikely to be composed of ryanodine receptors.

The lumenal material present within the SSC is intriguing; we are not aware of other examples of intracellular compartments with such regularly arranged structure in the lumen. This regularity is present in several ways: its location precisely along the mid-plane of the lumen, the hexagonal arrangement of discrete punctae, and its organization into parallel, equidistantly spaced rows. Because the central lumenal material is connected to its opposing membranes, an individual SSC cistern must be treated as a composite sandwich consisting of its two bounding membranes and the lumenal material. From a structural perspective, the SSC appears more akin to a truss bridge than the “floppy sack” that may be inferred from previous TEM studies that at times have revealed less uniform, empty cisterns likely due to fixation, extraction and dehydration artifacts typically associated with conventional sample preservation methods.

### Pillar Proteins Provide Thin Plasma Membrane-Cytoskeletal Connections

The wavy appearance of the PM in our preparations may be explained by the osmotic stress imposed by our HPF filler requirements, disrupting the regular spacing of density ascribed to pillar proteins in previous studies [24]. In cases of osmotic stress and the resulting PM variations, we find the pillar density joining PM to actin to vary in length and orientation, reflecting the variation in distance between the undulating PM and the actin filaments, whereas the actin filaments appear unaffected by the varying curvature of the PM and the resulting length differences of the pillar proteins, and instead remain tangent to the surface of the SSC membrane. This finding implies that the molecular interactions that connect the SSC to actin are stronger than those that connect the PM to actin.

Interestingly, regions of individual pillars are frequently as thin as ~3 nm and never exceed the cross-sectional area of an actin filament. The dimensions and the variable orientation raise the question of how such a delicate structure would effectively transmit the shear applied by a PM motor to the actin filament and avoid buckling in the process. Combined with the smooth appearance of the SSC membrane, its intermembrane quasi-periodic protein network, and preserved SSC-actin connections, it would seem that the pillar may be the weakest link in the coupling between PM, actin lattice, and SSC. This observation is consistent with micro-aspiration experiments demonstrating separation of the PM from the SSC-CL complex [44]. We therefore propose that the pillar protein mechanically functions more like an entropic spring rather than a rigid mechanical element. The recent labeling of the pillar with spectrin antibodies lends further support for this picture [45].

### Implications for OHC Mechanics

A cartoon schematic incorporating our findings into a new model of the lateral wall is illustrated in Figure 7a. At a minimum, the substructure visualized in the SSC could explain a cistern’s ability to maintain the spatial tolerances required of 9-10 μm diameter concentric cylinders of membranes spaced 20-30 nm apart and running the length of the OHC, spanning distances that can exceed 50 μm. Our structural findings are consistent with AFM studies indicating that the OHC possesses a highly elastic cortical shell organization [46]. This structure is also likely coupled to the maintenance of OHC turgor pressure, which is required for electromotility [47, 48]. The structural details of the SSC also have relevance for biophysical models of current flow within the cell and development of potential intracellular voltage gradients [49, 50]. The revealed molecular organization of the SSC raises the possibility that the SSC layer is mechanically anisotropic. In the radial direction, the lumenal material is densely packed and not likely to easily accommodate strain, whereas in the longitudinal direction, relative motion could potentially occur between the rows of densely packed material. Thus, the overall mechanical anisotropy of the outer hair cell lateral wall, previously identified solely with the actin-spectrin cytoskeleton [30], may also be reflected in the SSC. Previous results that the number of cisternae can vary depending on axial position along the OHC lateral wall, location in the cochlear spiral, and animal model [32, 38, 51], also raise the possibility that regulation of SSC structure may contribute to OHC specialization.

**Figure 7.**
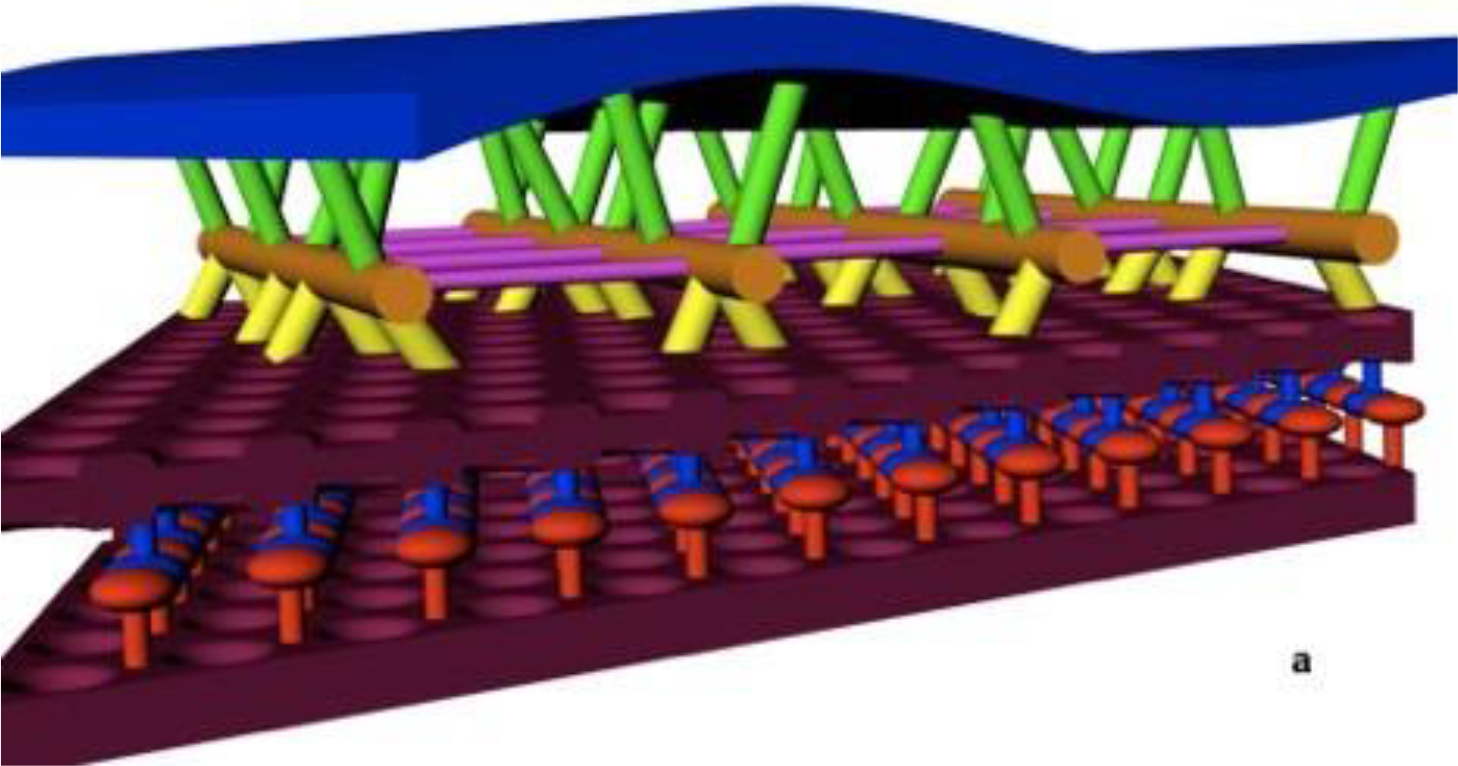
Model of the OHC lateral wall, including new SSC structure and actin-SSC links. Annotation: blue = PM; maroon = SSC membrane; orange = actin filament; yellow = actin-SSC link; green = pillar. For the SSC lumen material, red and blue are used to indicate the alternating material associated with each SSC membrane (see Figure 4i). Cross-links between actin, shown in other studies, are colored purple.

In summary, the novel ultra-structural findings presented here arise from a combination of advancements in sample preservation via HPF/FS, combined with the ability to resolve 3D structural relationships among components within the lateral wall through the use of electron tomography. The novel structural features observed motivate research to determine the molecular composition of the SSC membrane, the identity of cytoskeletal cross-linking molecules, and the implications of these findings for the biophysical mechanism of OHC electromotility.

## Materials and Methods

### Preparation of Samples

#### High-pressure freezing of OHCs

Hartley albino guinea pigs were anesthetized by isofluorane and decapitated with a guillotine. The temporal bones were removed and the organ of Corti was exposed by dissection in Invitrogen Medium 199 containing Hanks’ salts (Invitrogen Corp., CA, USA). Strips of the sensory epithelium were transferred using a 200 μL pipette and placed in a 35 mm MatTek dish (MatTek Corp, MA, USA); enzymatic digestion was avoided to preserve long strips of tissue. In brief, long strips containing OHCs were then identified by stereoscope and selected for further manipulation/handling using a microaspirator connected to 200 μm inner diameter Spectrapor dialysis tubing with a MW cutoff of 13-18kDa (Spectrum Labs, CA, USA). Using the microaspirator, samples were transferred to 10% glycerol (v/v), 20% dextran (w/v), or 20% BSA (w/v) as filler material and drawn back into the dialysis tubing. The tubing was then crimped to enclose the sample, transferred to either a 150 or 200 μm deep aluminum hat, the remaining space in the hat was filled with the same filler material, and the sample was cryoimmobilized using a BAL-TEC HPM-010 high-pressure freezer (BAL-TEC, Inc., Carlsbad, CA). Details of this procedure and the apparatus used are found in [36].

#### Freeze-substitution

Samples were freeze-substituted in 1% osmium tetroxide and 0.1% uranyl acetate in acetone using a Leica AFS (Leica Microsystems, Vienna, Austria) following a previously described protocol [52]. In some samples, 1-2% water was added to enhance membrane contrast [53]. Following freeze substitution, specimens were washed with several rinses of pure acetone before being infiltrated in either an Epon-Araldite mixture [54] or Durcupan ACM (Electron Microscopy Sciences, PA, USA). The tubes were flat-embedded between two slides using two layers of parafilm as a spacer [55].

#### Section preparation

Flat-embedded tubes were screened, mounted on blank epoxy blocks for sectioning, and their orientation for sectioning recorded using an Olympus SZX12 stereoscope (Olympus America, Inc., PA, USA) with epi- and trans-illumination provided by a pair of Fostec 150W lamps (Olympus). 150 nm sections were collected on formvar coated slot grids using a Reichert Ultracut E ultramicrotome (Leica). Sections were post-stained using a sequence of either 2% uranyl acetate in 70% methanol followed by Sato’s lead citrate[56], or 2% KMnO4 followed by a PALS bleach step and Sato’s lead citrate. 5 or 10 nm gold fiducials were added to both sides by floating on a drop of colloidal gold (British BioCell International, Cardiff, UK) following pre-treatment of each side with 0.001% (w/v) poly-L-lysine in double-distilled water.

### Electron tomography

All data presented represent dual-axis tilt series taken by rotation about orthogonal axes. Tilt series of 150 nm sections were collected at 1° increments through a range of +/− 70° using EMMENU 3.0 software (TVIPS GmbH, Gauting, Germany) on a Phillips CM200 TEM (FEI, Eindhoven, Netherlands) equipped with a Fischione Advanced Tomography Holder (E. A. Fischione Instruments, Inc., PA, USA) and a Tietz TEMCam-F214 2k × 2k CCD camera (TVIPS), or using SerialEM[57] on a JEOL JEM-2100F (JEOL Ltd,, Tokyo, Japan) equipped with a Tietz TEMCam-F415 4k × 4k CCD (TVIPS). Images were collected with an ~8 Angstrom pixel size at 1 – 1.5 μm defocus to preserve ~20 Angstrom information in the raw data, as evidenced by Thon ring position in resulting micrograph power spectra. Series were aligned with 5 or 10 nm gold fiducials and reconstructed using weighted back-projection with the IMOD software package [58]. Post-processing and segmentation of the resulting volume density maps was done using a combination of IMOD and the UCSF Chimera package [59, 60].

## Acknowledgements

We would like to thank Eva Nogales of UC Berkeley for usage of her Phillips CM200 TEM, and Henning Stahlberg of UC Davis for time on the JEOL JEM-2100F TEM in the Interdisciplinary Center for Electron Microscopy at UC Davis. We further would like to thank Dr. Patanjali Varanasi for fruitful discussions. This project was supported by a DOE CSGF fellowship administered by the Krell Institute, grant DE-FG02-97ER25308 (WJT), an NIH/NIDCD Ruth L. Kirschstein NRSA F30 MD/PhD award, grant 5F30DC009359 (WJT), an NSF CAREER Award, grant BES 044379 (RMR) and NIH/NIDCD grants DC009622 (RMR) and DC0008134 (RMR), NIH grant DC07680 (MA) and by the Director, Office of Science, of the U.S. Department of Energy, DE-AC03-76SF00098 (MA).

